# Theta Phase-Dependent Modulation of Perception by Concurrent tACS and Periodic Visual Stimulation

**DOI:** 10.1101/738906

**Authors:** Elif Somer, John Allen, Joseph Brooks, Vaughan Buttrill, Amir-Homayoun Javadi

## Abstract

**Background:** Sensory perception can be modulated by the phase of neural oscillations, especially in the theta and alpha ranges. Oscillatory activity in the visual cortex can be entrained by transcranial alternating current stimulation (tACS) as well as periodic visual stimulation (i.e., flicker). Combined tACS and visual flicker stimulation modulates blood-oxygen-level-dependent (BOLD) responses and concurrent 4 Hz auditory click-trains and tACS modulates auditory perception in a phase-dependent way.

**Objective:** In the present study, we investigated if phase synchrony between concurrent tACS and periodic visual stimulation (i.e., flicker) can modulate performance on a visual matching task.

**Methods:** Participants completed a visual matching task on a flickering visual stimulus while receiving either in-phase (0 degree) or asynchronous (180, 90, or 270 degrees) tACS at alpha or theta frequency. Stimulation was applied over either occipital cortex or dorsolateral prefrontal cortex (DLPFC).

**Results:** Visual performance was significantly better during theta frequency tACS over the visual cortex when it was in-phase (0 degree) with visual stimulus flicker, compared to anti-phase (180 degree). This effect did not appear with alpha frequency flicker or with DLPFC stimulation. Furthermore, a control sham group showed no effect. There were no significant performance differences amongst the asynchronous (180, 90, and 270 degrees) phase conditions.

**Conclusion:** Extending previous studies on visual and auditory perception, our results support a crucial role of oscillatory phase in sensory perception and demonstrate a behaviourally relevant combination of visual flicker and tACS. The spatial and frequency specificity of our results have implications for research on the functional organisation of perception.

## 1 Introduction

Previous research suggests that the state of oscillations in the brain may be a modulatory mechanism affecting sensory perception in both visual (e.g., Busch, Dubois, & VanRullen, 2009; Busch & VanRullen, 2010; Mathewson, Gratton, Fabiani, Beck, & Ro, 2009) and auditory domains (e.g., Lakatos et al., 2005; Ng, Schroeder, & Kayser, 2012; Rice & Hagstrom, 1989). In particular, perceptual performance can vary according to the phase of neural oscillations at the time of stimulus presentation (e.g., Mathewson, et al., 2009). Using brain stimulation techniques such as transcranial alternating current stimulation (tACS) to apply weak oscillatory electrical currents over the scalp (Kanai, Chaieb, Antal, Walsh, & Paulus, 2008: Thut, Schyns, & Gross, 2011), can modulate neural oscillations (Thut et al., 2011). Sensory stimulation, especially when periodic (i.e., flickering), can also affect neural oscillations. For instance, visual flicker stimulation can affect visual cognitive processes (e.g., Herbst, Javadi, van der Meer, & Busch, 2013: Herrmann, 2001). Steady state visual evoked potentials (SSEVP) are oscillatory neural responses to rhythmic visual stimuli observed in the electroencephalogram (EEG) at the flicker frequency (and its harmonics) primarily over the posterior scalp (e.g., Herbst, Javadi, van der Meer, & Busch, 2013; Herrmann, 2001; Spaak, de Lange, & Jensen, 2014; Brooks & Palmer, 2011). Further, both sensory and transcranial stimulation techniques were shown to be effective in phasic modulation of perception through entrainment of neural oscillations by EEG or magnetoencephalography (MEG) recordings and behavioural measures (Henry & Obleser, 2012; Jaegle, & Ro, 2014; Neuling, Rach, Wagner, Wolters, & Herrmann, 2012; Spaak et al., 2014).

Chai, Sheng, Bandettini, and Gao (2018) combined tACS with periodic visual stimulation. They reported blood-oxygen-level-dependent (BOLD) responses are modulated by tACS applied over Oz when it matched the visual flicker frequency or its second harmonic, mainly in areas activated by visual stimulation and targeted by the tACS current distribution. Further, Riecke, Formisano, Herrmann, and Sack (2015) combined near-threshold 4 Hz click trains and tACS applied over the auditory cortices with varying differences angles (30°, 90°, 150°, 210°, 270°, and 330) between the auditory stimulus and tACS waveform. They found that perceptual detection accuracy oscillated at the tACS frequency which is in line with other studies showing the phase of oscillations in the theta range is important for sensory perception (e.g. Busch et al., 2009; Busch & VanRullen, 2010; Lakatos et al., 2005; Ng et al., 2012; Tomassini, Ambrogioni, Medendorp, & Maris, 2017).

In the present study, we used concurrent tACS applied over the early visual cortex (V1) or dorsolateral prefrontal cortex (DLPFC) and visual stimulation in the theta (4.1 Hz) range to investigate if varying phase synchrony between them can affect perceptual performance in a visual matching task. We included DLPFC as a control location as it is associated with higher level cognitive processes such as short and long-term memory (e.g., Crowley, Bendor, & Javadi, 2019; Curtis & D’Esposito, 2003; Javadi, Glen, Halkiopoulos, Schulz, & Spiers, 2017; Javadi & Walsh, 2012) Therefore, it would not be likely to affect lower level perceptual performance. We also investigated if alpha stimulation (10 Hz) would affect task performance as this is another frequency range that has been repeatedly linked to visual perception (e.g., Babiloni, Vecchio, Bultrini, Luca Romani, & Rossini, 2005; Busch et al., 2009; Chai et al., 2018; Kanai et al., 2008; Mathewson et al., 2009; Spaak et al., 2014).

## 2 Materials and methods

### 2.1 Participants

The participants across all experiments were 105 University of Kent students (69 females, age range: 18-31), who received course credits or £8 as compensation for their time. They were screened for past and present neurological conditions and a number of health conditions that would prevent them from safely receiving electrical brain stimulation. They did not report any condition that could prevent them from participating in the study. All participants gave written informed consent. The experimental procedures were approved by the School of Psychology Ethics Committee at the University of Kent.

### 2.2 Stimuli & Apparatus

The experiment was presented on a 23” computer monitor (60 Hz, 1,920 × 1,080 pixel resolution) using the experiment software PsychoPy (Peirce, et al., 2019). The screen was situated at 60 cm viewing distance and the background colour was grey (RGB: 160, 160, 160). The visual display on each trial comprised four, vertically-oriented pseudo-random curvy lines (Figure 1) of 8.1° in height and a maximum of 3.8° in width. These were created using an algorithm from Brooks & Driver (2010). One target line appeared at the middle top of the display. Three curvy lines appeared in the bottom half of the display (Figure 1). One of these lines matched the target line whereas the other two were modified versions of the target line. The position of the matching line was random on each trial. Stimuli remained on the screen until response. The colour of the lines oscillated between magenta and the background grey (see subsection Visual stimulation and transcranial alternating current stimulation). Following response, masks appeared at the site of each stimulus. These comprised magenta random shapes on a grey background (Figure 2).

**Figure 1.**
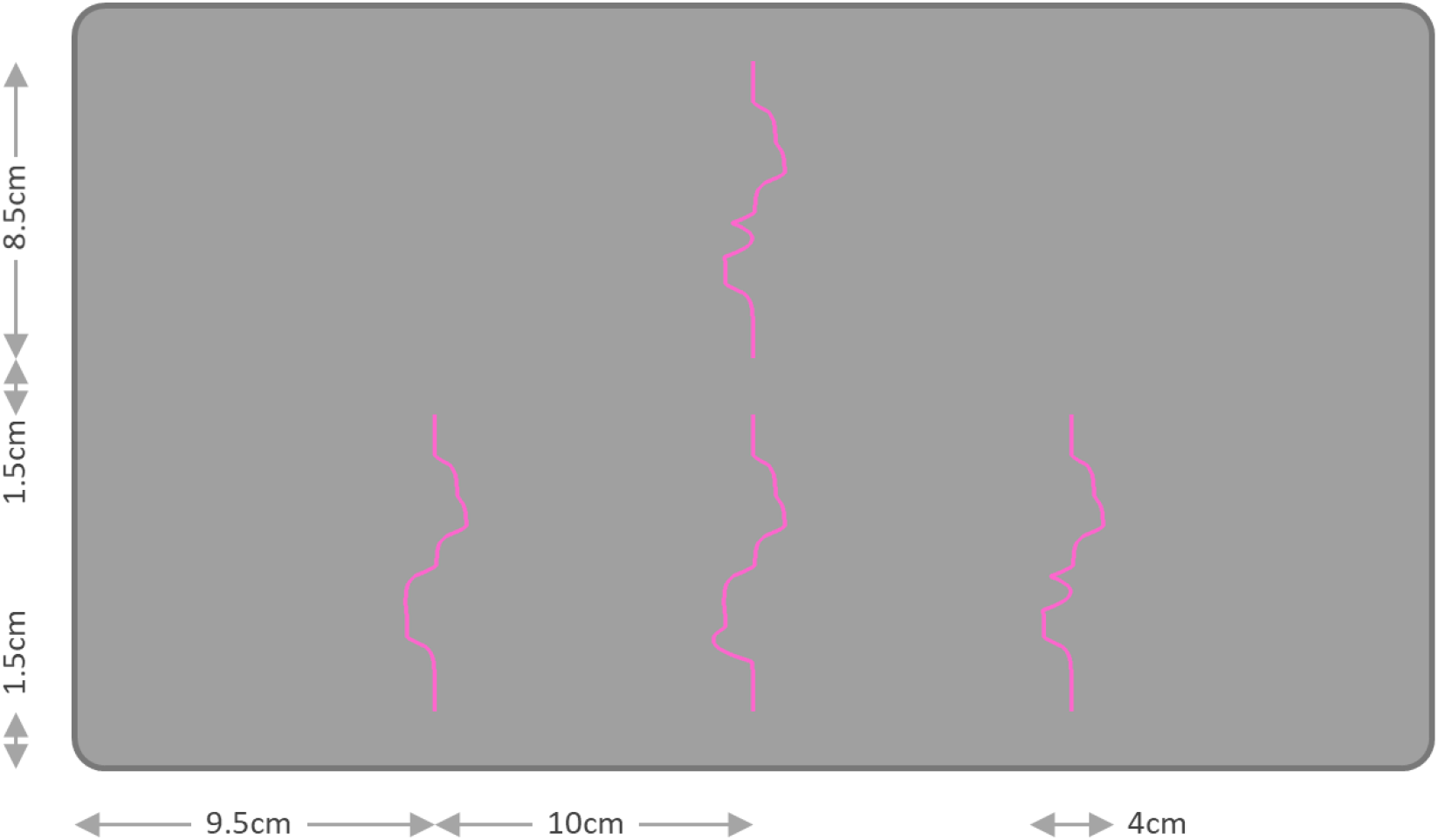
Static example of visual stimulus with the target edge on top and three match options along the bottom. In this example, the right-most stimulus on the bottom is the match to the target. The viewing distance was 60cm.

**Figure 2.**
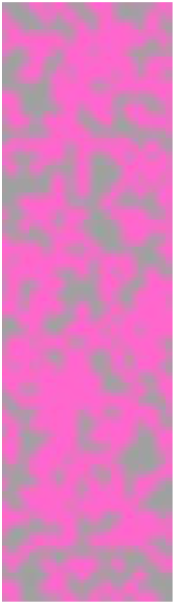
An example of the mask display.

### 2.3 Visual Stimulation and Transcranial Alternating Current Stimulation

The electrical brain stimulation was delivered using a battery operated NeuroConn stimulation device (Germany). Nuprep skin preparation gel and saline solution were used to improve conductivity. The area of stimulation was cleaned with skin preparation gel, and 5 × 7 cm2 rubber electrodes placed inside sponges soaked in saline solution were attached to the participants’ scalp and wrist using elastic bandages. The position of the electrodes attached to the scalp was defined by the 10-20 International EEG Electrode Placement System. For the participants in the occipital stimulation condition, the electrode was placed at position Oz. For the participants in the DLPC stimulation condition, the electrode was placed at position F3. The impedance was kept below 15 kΩ.

Alpha frequency stimulation was delivered at 10 Hz and theta frequency stimulation was delivered at 4.1 Hz depending on the condition the participant was randomly assigned to. The stimulation waveform always started at zero amps (i.e., zero phase). To create visual flicker, the colour of the stimuli oscillated between magenta and the background grey at either 4.1 Hz or 10 Hz. Visual stimuli onset was at 0°, 90°, 180°, or 270° phase relative to the tACS waveform. For instance, in the 0° phase condition, the visual stimulus would start to ramp up towards full visibility at the 0° position (zero amps ascending) of the tACS waveform. See the next section for details on how phase conditions were combined across experiments.

### 2.4 Design & Procedure

The first block of 130 trials were practice trials. No stimulation was applied in this block. Next, during the two stimulation blocks, a 4.1 or 10 Hz (depending on the experiment) sinusoidal current was applied at every trial. The stimulation was ramped up to 1.5 mA over four cycles at the beginning of each mini-block. Mini-blocks lasted for either 10 trials or 30 seconds, whichever was shorter depending on the response speed of the participant. Visual stimuli oscillated (starting as the background colour) throughout the task at the same frequency as tACS in that experiment. The phase of the onset of the visual stimuli relative to tACS was randomly selected on each trial. The phases used varied across experiments (see below). Mini-blocks were separated by 8 seconds. There were a total of 130 trials in each of the two stimulation blocks.

On each trial, participants performed a visual matching task. The participants were instructed to indicate which of three curvy lines matched the target using the left, down, or right arrow keys on a keyboard. Each trial started with a 650-ms fixation cross followed by the stimuli. The stimuli stayed on the screen until the participant responded. After the participant responded, masks (see Stimuli & Apparatus) were presented for 100 ms, followed by the next trial. Inter-trial interval varied as a result of reaction time in the trial.

In Experiment 1, we used a 2 × 2 mixed-factors design with 30 participants (15 in each stimulation location) completed a visual matching task with a theta frequency flickering visual stimulus while receiving tACS, at the same frequency, over either the occipital cortex or DLPFC. We implemented a 2 × 2 mixed factorial design. The factors were stimulation phase (0°, 180°; within-subjects) and stimulation location (DLPFC or occipital; between-subjects).

In Experiment 2, in order to investigate if the observed effect would vary with asynchronous phase delays other than the anti-phase (180°), we tested 30 more participants with 90° and 270° phase difference between visual stimulation and tACS.

In Experiment 3, we repeated Experiment 1 with 30 new participants in order to investigate if the effect we found can be observed with stimulation in the alpha range (10 Hz). All other aspects of the study were kept the same.

In Experiment 4, we ran a control experiment using in- or anti-phase labels randomly assigned to trials and occipital or DLPFC electrode placements without actual stimulation with 15 participants.

### 2.5 Statistical analysis

The participants’ reaction times and accuracy at every trial were recorded. The inverse efficiency score (IES, mean reaction time by the proportion of correct responses) (Bruyer, & Brysbaert, 2011; Townsend, & Ashby, 1978), was calculated and used in the subsequent analyses. We used an alpha level of .05 for all statistical tests. The assumption of homogeneity of variance were met (Levene’s tests, *p* > .05) for all of the statistical analyses unless stated otherwise. There were violations of the normality assumption (Shapiro-Wilk tests, *p* < .05) in some of the conditions. However, the group sizes were equal (or almost equal in Experiment 4) and no simple main effects tests were conducted for these groups.

## 3 Results

### 3.1 Experiment 1 – modulation of perception by phase synchrony

The results of Experiment 1 showed that differences in phase synchrony between visual stimulation and tACS affected perceptual performance. A mixed design 2 × 2 ANOVA with stimulation phase (0°, 180°) as a within-subjects factor, stimulation location (occipital, DLPFC) as a between-subjects factor, and task performance (Inverse Efficiency Scores [IES]: see Methods) as the dependent variable was conducted. There was a significant interaction between stimulation phase and location, F(1, 27) = 4.72, *p* =.038, *η_p_^2^* = .14 (see Figure 3), but no significant main effect of stimulation phase, F(1, 27) = 2.72, *p* = .110, *η_p_^2^* = .09, or location, F(1, 27) = .29, *p* = .596, *η_p_^2^* = .01. Pairwise comparisons with Bonferroni corrections revealed that performance was better in the 0° compared to 180° phase condition in the occipital group, t(14) = 2.70, *p* = .012, d = 0.70, but there was no significant difference for the DLPFC group, t(14) = .37, *p* = .714, d = 0.09 (See Table 1 for the descriptive statistics and estimated mean and standard error values). Levene’s test of equality of error variances was significant, *p* = .022, for the 180° phase condition. However, the main effect of the between subjects factor and pairwise comparisons between occipital and DLPFC groups for both 180°, t(14) = .14, *p* = .889, d = 0.04, and 0°, t(14) = .98, *p* = .334, d = 0.25, conditions were already not significant.

**Figure 3.**
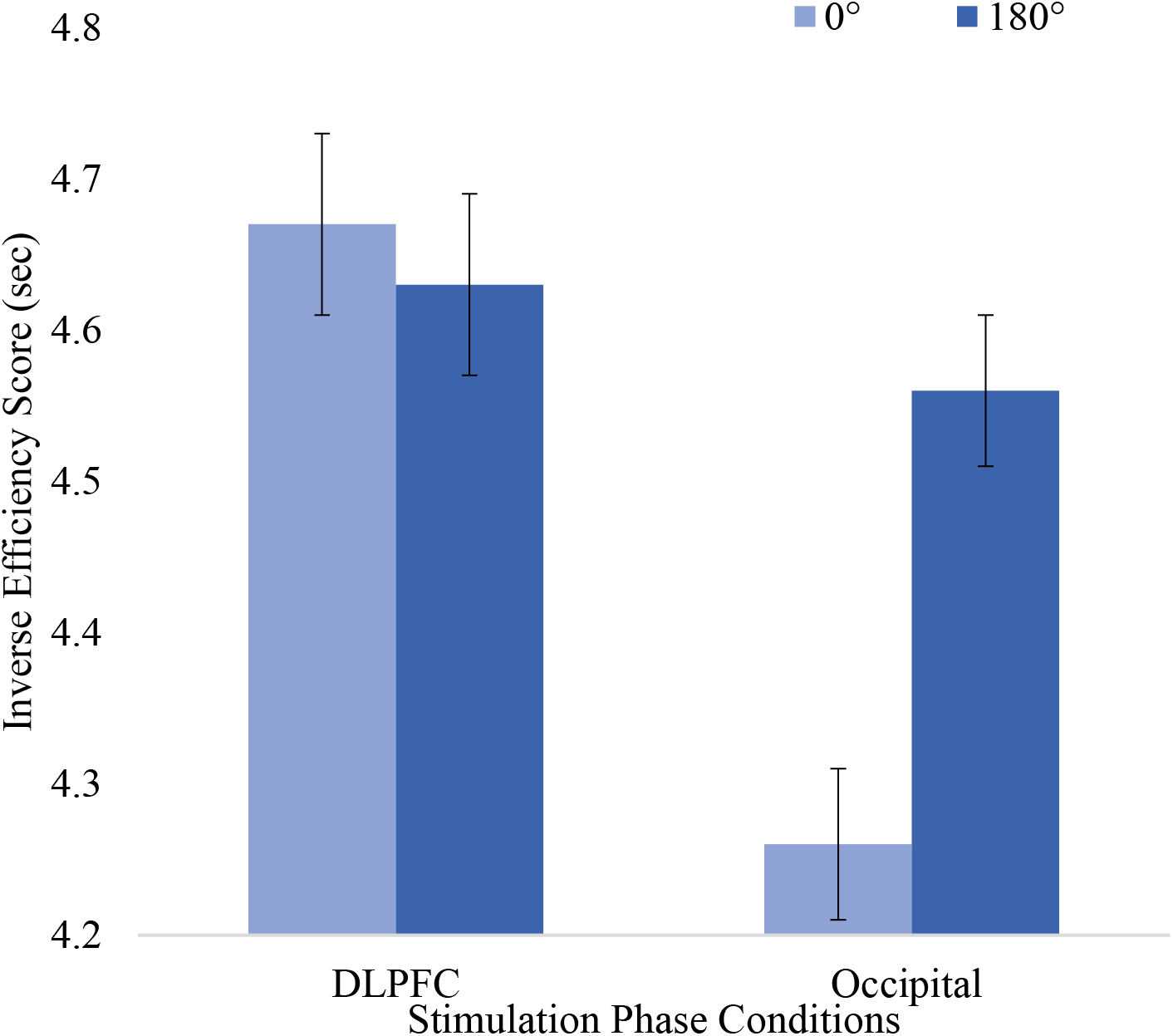
Estimated Marginal Means of inverse efficiency scores (IES) in Stimulation Phase Conditions by Stimulation Location Groups in Experiment 1. Error bars represent one standard error adjusted for related means

**Table 1.** Descriptive Statistics and estimated mean and standard error values of the inverse-efficiency scores (IES, (mean[SD]) for stimulation phase conditions according to stimulation location groups.

| Stim. Location | Degrees | Exp. 1 | Exp. 2 | Exp. 3 | Exp. 4 |
| --- | --- | --- | --- | --- | --- |
| Occipital | 0° | 4.26[1.33] | 4.13[0.91] | 4.41[1.60] | 3.99[1.61] |
|  | 180° | 4.56[1.64] | 4.07[0.89] | 4.45[1.68] | 4.10[1.80] |
| Est. Occipital | 0° | 4.26[1.49] | 4.13[1.49] | 4.34[0.87] | 3.99[2.46] |
|  | 180° | 4.56[1.74] | 4.07[1.43] | 4.38[0.82] | 4.10[2.66] |
| DLPFC | 0° | 4.67[0.93] | 4.05[1.27] | 4.33[1.15] | 4.04[0.87] |
|  | 180° | 4.63[0.92] | 4.20[1.23] | 4.16[1.09] | 4.05[0.85] |
| Est. DLPFC | 0° | 4.26[1.33] | 4.13[0.91] | 4.41[1.60] | 3.99[1.61] |
|  | 180° | 4.56[1.64] | 4.07[0.89] | 4.45[1.68] | 4.10[1.80] |

Notes: Stim.: Stimulation, Est.: Estimated

As we used a combined measure of RT and accuracy for our main analyses, we conducted additional analyses to investigate the source of the phase synchrony effect further. Our results indicate that the observed effect was mainly driven by RT rather than accuracy. A mixed design 2 × 2 ANOVA was repeated with stimulation phase (0°, 180°) as a within-subjects factor, stimulation location (occipital, DLPFC) as a between-subjects factor, and task performance (RT) as the dependent variable. There was a significant interaction effect between stimulation phase and location, F(1, 27) = 7.82, *p* =.009, *ηp^2^* = .22 (see Figure 4), but no significant main effect of stimulation phase, F(1, 27) = 2.92, *p* = .099, *η_p_^2^* = .09, or location, F(1, 27) = .26, *p* = .616, *η_p_^2^* = .01. Performance was worse in the 180° compared to 0° phase condition in the occipital group, t(14) = 3.20, *p* = .004, d = 0.83, and there was again no significant difference for the DLPFC group, t(14) = .77, *p* = .448, d = 0.20, as shown by pairwise comparisons with Bonferroni corrections (See Table 2 for the descriptive statistics and estimated mean and standard error values). Levene’s test of equality of error variances was significant, *p* = .014, for the 180° phase condition. However, the main effect of the between subjects factor and pairwise comparisons between occipital and DLPFC groups for both 180°, t(14) = .16, *p* = .873, d = 0.04, and 0°, t(14) = .88, *p* = .386, d = 0.23, conditions were already not significant.

**Figure 4.**
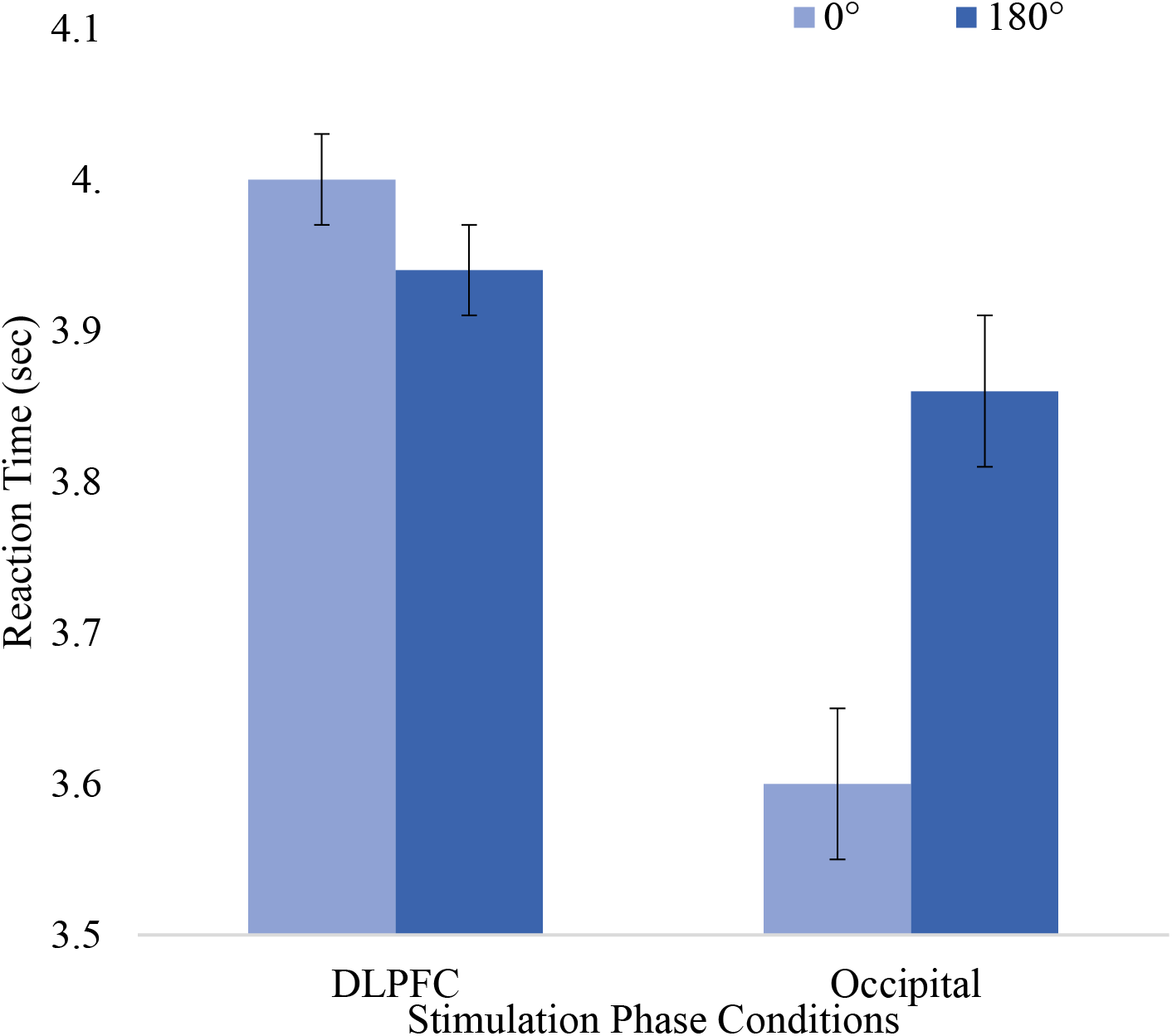
Estimated Marginal Means of RTs in Stimulation Phase Conditions by Stimulation Location Groups in Experiment 1. Error bars represent one standard error adjusted for related means

**Table 2.** Descriptive Statistics and estimated mean and standard error values of the, reaction times (RT) and proportion of correct responses (PC, mean[SD]) for stimulation phase conditions according to stimulation location groups in Experiment 1.

| Stimulation Location | Degrees | RT | PC |
| --- | --- | --- | --- |
| Occipital | 0° | 3.60[1.40] | 0.83[0.15] |
|  | 180° | 3.86[1.67] | 0.82[0.14] |
| Est. Occipital | 0° | 3.60[1.69] | 0.83[0.15] |
|  | 180° | 3.86[1.84] | 0.82[0.15] |
| DLPFC | 0° | 4.01[1.11] | 0.85[0.10] |
|  | 180° | 3.94[1.04] | 0.85[0.12] |
| Estimated DLPFC | 0° | 4.01[1.69] | 0.85[0.15] |
|  | 180° | 3.94[1.84] | 0.85[0.15] |

A mixed design 2 × 2 ANOVA with stimulation phase (0°, 180°) as a within-subjects factor, stimulation location (occipital, DLPFC) as a between-subjects factor, and task performance (proportion of correct responses [PC]) as the dependent variable revealed no significant interaction effect between stimulation phase and location, F(1, 27) = .09, *p* =.773, *η_p_^2^* = .00, and no significant main effect of stimulation phase, F(1, 27) = .05, *p* = .824, *η_p_^2^* = .00, or location, F(1, 27) = .28, *p* = .603, *η_p_^2^* = .01 (See Table 2 for the descriptive statistics and estimated mean and standard error values).

### 3.2 Experiment 2 – no significant difference between ansynchronous phase conditions

In Experiment 2, a mixed design 2 × 2 ANOVA with stimulation phase (90°, 270°) as a within-subjects factor, stimulation location (occipital, DLPFC) as a between-subjects factor, and task performance (IES) as the dependent variable revealed no significant interaction effect between stimulation phase and location, F(1, 27) = 2.92, *p* =.098, *η_p_^2^* = .10, and no significant main effect of stimulation phase, F(1, 27) = .70, *p* = .410, *η_p_^2^* = .02, or location, F(1, 27) = .01, *p* = .946, *η_p_^2^* = .00 (See Table 1 for the descriptive statistics and estimated mean and standard error values). Four independent-samples t-tests between the asynchronous phase conditions from Experiments 1 and 2 for the occipital and DLPFC groups were conducted. As Levene’s tests indicated the assumption of equality of variances was violated, the degrees of freedom for the occipital group were adjusted. There were no significant differences between phases 180° and 90°, or 270° in the occipital group. There were also no significant differences between phases 180° and 90°, or 270 in the DLPFC group (see Table 1 for mean and standard deviation values, and Table 3 for a summary of the independent samples t-tests).

**Table 3.** Summary of the Independent Samples T-tests Comparing Phase Asynchronous Conditions from Experiments 1 and 2.

| Stimulation Location | Stim. Phase | t | p | Cohen's d |
| --- | --- | --- | --- | --- |
| Occipital | $180^\circ - 90^\circ$ | $t(21.80)=0.90$ | .378 | 0.33 |
| | $180^\circ - 270^\circ$ | $t(21.61)=2.69$ | .323 | 0.37 |
| DLPFC | $180^\circ - 90^\circ$ | $t(28)=1.43$ | .164 | 0.52 |
| | $180^\circ - 270^\circ$ | $t(28)=1.08$ | .290 | 0.39 |

### 3.3 Experiment 3 – frequency specificity of effects

In Experiment 3, there was no significant interaction effect between stimulation phase and location, F(1, 27) = 2.70, *p* =.111, *η_p_^2^* = .09, and no significant main effect of stimulation phase, F(1, 27) = 1.07, *p* = .310, *η_p_^2^* = .04, or location, F(1, 27) = .13, *p* = .720, *η_p_^2^* = .01 in a mixed design 2 × 2 ANOVA with stimulation phase (0°, 180°) as a within-subjects factor, stimulation location (occipital, DLPFC) as a between-subjects factor, and task performance (IES) as the dependent variable (See Table 1 for the descriptive statistics and estimated mean and standard error values).

### 3.4 Experiment 4 – no effect of Sham-tACS

Finally in Experiment 4, there was no significant interaction, F(1, 13) = .34, *p* = .571, *η_p_^2^* = .03, or main effects of assigned phase, F(1, 13) = .41, *p* =.531, *η_p_^2^* = .03, or location, F(1, 13) = .00, *p* =.999, *η_p_^2^* < .001, in a mixed design 2 × 2 ANOVA with assigned phase (0°, 180°) as a within-subjects factor, electrode location (occipital, DLPFC) as a between-subjects factor, and task performance (IES) as the dependent variable (See Table 1 for the descriptive statistics and estimated mean and standard error values).

## 4 Discussion

We investigated the effects of varying phase synchrony between concurrent theta (4.1 Hz) or alpha (10 Hz) tACS and visual flicker applied over the occipital cortex or DLPFC. Our findings showed that phase synchrony between theta visual stimulation and tACS applied over the visual cortex modulates perceptual performance. Performance was significantly better in the in-phase (0°) stimulation condition compared to anti-phase (180°) stimulation condition only in the theta occipital group. Our results are largely in line with Riecke et al. (2015) who showed phase synchrony between 4 Hz click trains and tACS modulated auditory perception. However, they found performance was better for the stimuli presented during the positive half-wave (phases 30° and 150°) of 4-Hz tACS than the negative half-wave (phases 210° – 330°) while we found no difference between asynchronous phase conditions. Clouter, Shapiro, and Hanslmayr (2017) who showed better performance with phase synchrony between oscillating theta auditory and visual stimuli also reported no difference between varying asynchronous phase offsets. However, their outcome measure was episodic memory. Our findings are unlikely to be due to a general tACS effect on motor performance as the phase of tACS relative to the visual stimulus would not be expected to affect performance in such case. Further, electrophysiological evidence for the efficiency of tACS applied over the occipital cortex in phase-specific modulation of SSVEPs has recently been reported, although in the alpha frequency range (Fiene et al., 2019). More broadly, our findings support the literature suggesting sensory perception is modulated by the phase of oscillations in the sensory cortices (e.g. Busch et al., 2009; Busch & VanRullen, 2010; Lakatos et al., 2005; Mathewson et al., 2009; Neuling et al., 2012; Ng et al., 2012; Rice & Hagstrom, 1989; Riecke et al, 2015) and the importance of theta range oscillations for perception (e.g. Busch et al., 2009; Busch & VanRullen, 2010; Lakatos et al., 2005; Ng et al., 2012; Riecke et al, 2015; Tomassini et al., 2017).

Interestingly, the effect that we observed during theta stimulation was not present for alpha stimulation. Both theta (e.g. Demiralp et al., 2007; Lakatos et al., 2005; Landau, Schreyer, Van Pelt, & Fries, 2015; Spyropoulos, Bosman, & Fries, 2018) and alpha (e.g. Herring, Esterer, Marshall, Jensen, & Bergmann, 2019; Spaak, Bonnefond, Maier, Leopold, & Jensen, 2012) oscillations have been associated with the modulation of visual processing through modulating gamma oscillations. However, alpha oscillations were associated with selective attention on a single stimulus (e.g. Thut, Nietzel, Brandt, & Pascual-Leone, 2006) and suppressing information from unattended regions (e.g. Händel, Haarmeier, & Jensen, 2011). while distributed attention between two or more stimuli is facilitated by theta rhythmicity (e.g. Fiebelkorn, Saalmann, & Kastner, 2013; Kienitz et al., 2018; Landau, & Fries, 2012; Landau et al., 2015; Spyropoulos et al., 2018). Our task required comparing at least two stimuli simultaneously while most studies focusing on alpha oscillations included perception of a single, often near-threshold stimulus at a time (e.g. Babiloni et al., 2005; Busch et al., 2009; Mathewson et al., 2009; Rice & Hagstrom, 1989; Spaak et al., 2014). The need to investigate possible differences in the functions of alpha and theta frequency bands for the perceptual process depending on the task or stimuli has been previously discussed (Hanslmayr, Volberg, Wimber, Dalal, & Greenlee, 2013). Our results indicate differential effects of the two frequency bands on perception. Future research can address if with a single near-threshold stimuli detection task the effect would be observed in the alpha range, although this could present challenges to delivering effective visual stimulation. Further, our effect mainly stemmed from reaction time rather than accuracy. Distinct effects of attentional cueing characteristics on reaction time and accuracy have previously been shown, and they may represent different underlying processes (e.g. Prinzmetal, McCool, & Park, 2005; van Ede, de Lange, & Maris, 2012). However, further research would be needed to determine if our task characteristics interact with these two measures differently.

As well as modulating visual attention and perception within the visual cortex by facilitating bidirectional communication and influence between lower and higher levels (Hanslmayr et al., 2013; Spyropoulos et al., 2018) neural oscillatory activity is a potential mechanism for dynamic organisation and communication of cognitive processes of all levels across modalities and distant brain areas (Başar, Başar-Eroğlu, Karakaş, & Schürmann, 2000, 2001; Buzsáki, & Draguhn, 2004; Engel, Fries, & Singer, 2001) particularly in the theta range (Başar et al., 2000, 2001; Demiralp et al., 2007; Hanslmayr et al., 2013; Spyropoulos et al., 2018). It remains an open research question how the demonstrated effect would be reflected in later cognitive processes such as memory and what the effects of varying phase synchrony between sensory stimulation and tACS applied over parietal or frontal regions be on memory. This presents an opportunity for investigating the functional and anatomical structure of perceptual and memory processes and the relationship between them. Future research can also employ simultaneous tACS, visual stimulation and magnetoencephalographic (MEG) recording using a recently demonstrated novel approach (Ruhnau, Keitel, Lithari, Weisz, & Neuling, 2016). Although SSEVPs are well documented (e.g. Herbst et al., 2013; Herrmann, 2001; Spaak et al., 2014). and there is emerging evidence of entrainment through tACS (e.g. Helfrich et al., 2014; Kanai et al., 2008), electrophysiological evidence would strengthen and further clarify our findings as discussed by Thut et al. (2011).

Our Study contributes to the recently developing literature on concurrent use of tACS and periodic visual stimulation as a promising research technique in neuroscience (Ruhnau et al, 2016: Chai et al., 2018) and the importance of phase synchrony between them (Fiene et al., 2019) and demonstrate a behavioural effect for the first time. This will be of interest to researchers using electrical brain stimulation and sensory stimulation techniques. It might also have implications for applied researchers focusing on the therapeutic and performance enhancement uses of electrical brain stimulation. Further, the spatial, frequency, and phase specificity of our findings would be significant for all researchers interested in the biological and functional organisation of perception.

